# Early life stress programming of NG2+ glia transcriptome alters functional properties of voltage gated sodium (Nav) channels and cognitive performance

**DOI:** 10.1101/2020.08.19.257113

**Authors:** Giulia Treccani, Hatice Yigit, Thomas Lingner, Vanessa Schleuβner, Malin Wennström, David P Herzog, Markus Fricke, Gregers Wegener, Thomas Mittmann, Jacqueline Trotter, Marianne B Müller

## Abstract

The precise mechanisms underlying the detrimental effects of early life stress (ELS) on adult mental health remain still elusive. To date, most studies have exclusively targeted neuronal populations and not considered neuron-glia crosstalk as a crucially important element for the integrity of stress-related brain function. Here, we have investigated the impact of ELS on a glial subpopulation with unique properties in brain homeostasis, the NG2+ cells. ELS shifted the NG2+ transcriptome towards more mature stages, and these transcriptional effects were dependent on stress-induced glucocorticoids. The functional relevance of one candidate gene, Scn7a, could be confirmed by an increase in the density of voltage-gated sodium (Nav) channel activated currents in hippocampal NG2+ cells. Scn7a remained upregulated until adulthood in ELS animals, and these same animals displayed impaired cognitive performance. Considering that Nav channels are important for NG2+ cell-to-neuron communication, our findings suggest novel insights into the pathophysiology of stress-related mental disorders.

## Introduction

The brain is particularly sensitive to adversity experienced early in life (1). Early adversity (EA)-such as child neglect and maltreatment-lastingly impacts developmental trajectories of the brain, leading to structural changes (e.g. reduced cortical and hippocampal volumes) but also functional impairment of critical neurocognitive circuits in adulthood (2–4). In addition, meta-analytical data confirm that exposure to EA during childhood not only increases the risk for developing neuropsychiatric disorders later in life but also predicts an unfavorable course of illness and treatment outcome in depression (5–7). So far, the vast majority of studies into the neurobiology of ELS have exclusively targeted alterations in neuronal function. In recent years, the importance of neuron-glia communication and bidirectional neuron-glia crosstalk is emerging as crucial for proper brain function throughout the lifespan and has been implicated in brain disease-related processes(8). There is evidence that, e.g. impaired brain myelination contributes to the detrimental effects of EA on brain development across species (9, 10), but knowledge about the precise molecular mechanisms and glial subpopulations involved is still lacking. Despite its remarkable potential to adapt to different extrinsic stimuli and promote functional recovery (11, 12), myelination can be disturbed if environmental challenges are severe or long-lasting, e.g. under conditions of critical stress, which has been proven by both clinical (13) and preclinical studies (14, 15). Myelinating cells - oligodendrocytes within the central nervous system-derive from the differentiation of oligodendrocyte precursor cells (OPCs) (16). Both OPCs and mature oligodendrocytes can be affected by early stressful experiences (17, 18). OPCs expressing the type I proteoglycan CSPG4 (NG2) on their cell surface, thus known as NG2+ cells, have been discussed to play additional roles in brain homeostasis, although the exact nature of such roles remains unclear(19–22). In addition to serving as progenitors of myelinating oligodendrocytes (19, 23), NG2+ cells have unique properties, rendering them particularly interesting in the context of stress and EA. These properties include the ability to establish synaptic contacts with neurons (24–27), to communicate with microglia (28) and to modulate glutamatergic signal transduction in adjacent neurons via the shed version of the NG2 ectodomain (29). NG2+ cells express the glucocorticoid receptor (GR) (30) and their proliferation is regulated by glucocorticoids (31–33). Despite the interesting roles of NG2+ cells and their responsiveness to glucocorticoids (31, 32) and chronic stress in adulthood (34), so far no study has addressed the question as to whether NG2+ cells could be a target population of ELS. Similarly, it has not been addressed whether they could be involved in mediating the long-term negative consequences of ELS on mental health. In rodent models, ELS takes place during the so-called stress hypo-responsive period (SHRP) (35) which is a period characterized by a markedly reduced responsiveness of the hypothalamus-pituitary adrenal system to moderately challenging conditions. SHRP has been discussed to protect the early postnatal and still developing brain from an excess of glucocorticoid hormones (36). Intriguingly SHRP (postnatal day 2-9) largely overlaps with the temporal window during which NG2+ cells reach their peak density (the first postnatal week), and when the majority of newly divided NG2+ cells differentiate into oligodendrocytes (37). ELS is capable of overriding SHRP and eliciting a significant stress response with increased expression of central corticotropin releasing hormone and enhanced circulating corticosterone (CORT) concentrations, which in turn cause detrimental long lasting effect on memory formation and hippocampus-dependent cognitive functions (38–40). However, the individual contribution of an ELS-induced CORT excess and the concomitant activation of central stress neurocircuits on the developing brain remains to be fully dissected.

To this end, we used an established mouse model applying ELS through fragmented maternal care (41, 42) and performed cell-type specific transcriptional profiling in hippocampal NG2+ cells directly after stress exposure. To dissect the impact of CORT on modulating ELS-induced negative outcomes, we further correlated the molecular changes with the individual litter’s CORT response to ELS. Second, we performed enrichment analyses to test the hypothesis that glucocorticoid responsive genes are overrepresented in our set of ELS candidate genes and to determine whether ELS might shift the molecular identity/profile of NG2+ cells and alter the maturation stage within the lineage. Extending our analyses to the adult stage we identified an impairment of hippocampus-dependent cognitive performance to be accompanied by persistent changes in the NG2 transcriptome profile of adult animals previously exposed to ELS, thus suggesting *Scn7a* (Sodium channel protein type 7 subunit alpha) as a strong ELS candidate gene. Finally, the functional relevance of an ELS-induced increase in the expression of *Scn7a* was confirmed by electrophysiological recordings in hippocampal NG2+ cells.

## Materials and Methods

### Animals

Adult female and male C57bL/6J mice (10 weeks) were obtained from Janvier Labs (France). Mice were single housed with food and water ad libitum in an air-conditioned (temperature=22 ± 2 °C, relative humidity=50 ± 5%) housing room with 12 h/12 h light-dark cycle (lights on at 07:00 a.m.). Mice were habituated to the new environment for at least 1 week before the starting of the experiment. The breeding pairs consisted of female and male C57BL/6J. After mating pregnant dams were single housed and put in a cabinet with controlled ventilation, temperature and light cycle until the end of the experiment. From postnatal day (P) 2 to P9 the dams and litter were exposed to early life stress procedure (ELS) or control condition. For electrophysiological experiments C57bL/6N mice were bred to homozygous knock-in NG2 enhanced yellow fluorescent protein (NG2-EYFP) mice, where the expression of the reporter gene is regulated according to the endogenous NG2 promoter, allowing unbiased sampling of this population (43), to obtain heterozygous NG2-YFP pups. All experiments were performed in accordance with the European directive 2010/63/EU for animal experiments and were approved by the local authorities (Animal Protection Committee of the State Government, Landesuntersuchungsamt Rheinland-Pfalz, Koblenz, Germany).

### Early life stress protocol

ELS was performed as previously described (41, 42). Briefly, on P2, dams with 5-6 pups (of which at least one female) were randomly subjected to ELS or assigned to the control group from P2 to P9. The dams subjected to ELS were housed in cages with limited bedding and nesting materials (half square of a Nestlet, # 14010 Plexx) on an aluminum mesh platform as for (41)(McNichols Co) at least 1.5 cm above the cage floor. Control litters were kept under standard conditions and provided with sufficient bedding and nesting materials (2 Nestlets) All animals were monitored but left undisturbed until the end of the experiment. For the acute effect of ELS, in the morning of P9 the male pups were weighed and euthanized. For the long-lasting effect of ELS at P9 animals were weighed and were kept and housed again under standard conditions until weaning at P21. After weaning the male offspring were kept group housed until 8 weeks. At week 9-10 the animals were single housed, behavioral phenotype tested at 4-5 months and mice were euthanized at 8 months. For a schematic overview of the different experimental schedules see Fig 1.

**Fig 1:**
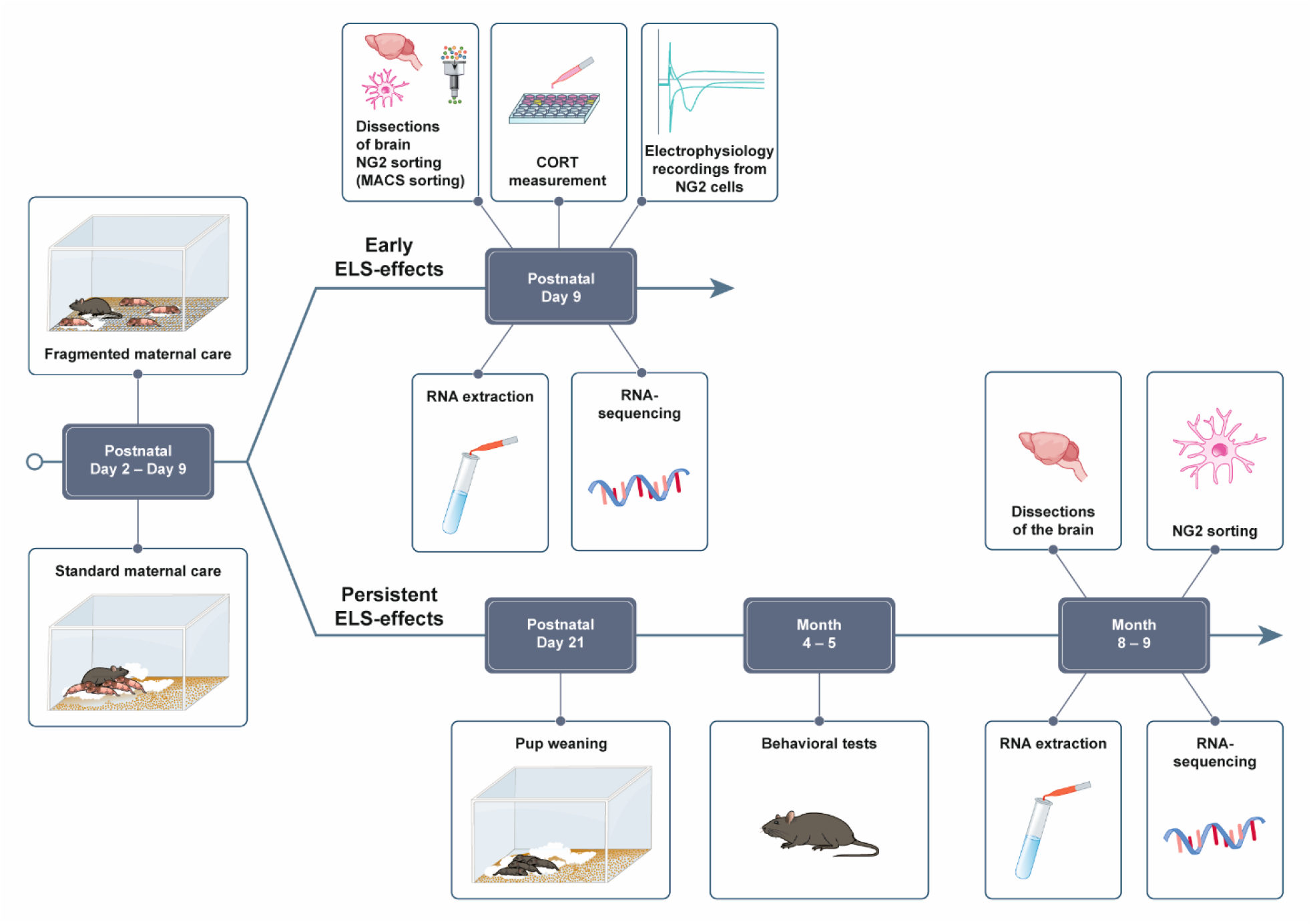
Schematic overview of the experimental procedures.

### Measurement of plasma CORT

Concentrations of circulating corticosterone were measured from trunk blood plasma (P9) or tail cut (adult animals) by Corticosterone ELISA Kit (Enzo Life Sciences, #Cat.No. ADI-900-0979) according to the manufacturer’s instructions. The blood was collected in an EDTA tube and centrifuged for 10 min at 4 °C and 10 000 rpm.

### Behavioral tests

All behavioral experiments were conducted between 09:00 and 14:00 by experimenters blinded to the treatment groups. All the animals were randomized per condition. The videos of all behavioral tests were scored manually by an experimenter blinded to the animals’ treatment using The Observer XT12 software (Noldus Information Technology). Between each test session, setups and arenas were cleaned with a 5% EtOH solution and dried with tissues. Behavioral testing occurred in custom-made sound-attenuating boxes under constant light conditions (37 lx). Detailed descriptions of the behavioral tests performed are provided in the Supplementary Information (SI).

### Isolation of NG2+ cells by magnetic cell sorting

NG2+ cells were isolated using the NTDK-P Kit (Miltenyi Biotec) according to previously described procedures (44, 45), where Magnetic isolation (MACS) was performed with anti-NG2 antibody conjugated with magnetic beads (Miltenyi Biotec). For the P9 time-point, each sample was a pool of the hippocampi from the male pups of the entire litter; n represents litter number (control n = 7, ELS low CORT n= 8, ELS high CORT n= 7, randomized in 3 batches). For the 8-month time-point each sample was a pool of the hippocampi of 3-4 males per sample (control n = 6, ELS n = 7 randomized in 3 batches).

### RNA extraction and Next-Generation Sequencing (NGS)

After NG2 sorting, cells were pelleted and RNA was extracted using RNeasy Micro Kit (QIAGEN) according to manufacturer’s instructions. NGS library prep was performed with NuGen Ovation SoLo RNA-Sequencing (RNA-seq) System following NuGen`s standard protocol (M01406v2). For the first experiment (at P9) libraries were prepared with a starting amount of 1 ng and amplified in 14 PCR cycles. Libraries were profiled in a High Sensitivity DNA on a 2100 Bioanalyzer (Agilent technologies) and quantified using the ddPCR Library Quantification Kit for Illumina TruSeq in a QX200 Droplet Digital PCR system (BioRad). Details information on the NGS protocol are described in the SI.

### Bioinformatic analysis

Details descriptions of the bioinformatic analysis are provided in the SI.

### Electrophysiology

Hippocampal NG2+ cells were electrophysiologically characterized at the age of P9-P10 (n=11 (control), n=11 (ELS)) by use of heterozygous knock-in mice expressing the enhanced yellow fluorescent protein EYFP in NG2+ cells (NG2-EYFP+/-mice,(43)). After decapitation the brains were quickly removed and put into ice-cold 95% O_2_/5% CO_2_ oxygenated ACSF. Next, the tissue was horizontally cut by use of a vibratome (VT 1200S, LEICA, Germany) to generate hippocampal brain slices with a thickness of 300 μm. The slices were kept in oxygenated ACSF at room temperature for at least 1 hour for recovery. Then they were transferred into a submerged-type recording chamber mounted on an upright microscope (Olympus BX-51WI) and perfused with oxygenated 95% O_2_/5% CO_2_ HEPES-buffered recording solution at room temperature for 10 minutes. NG2+ cells located in the stratum radiatum of area CA1 could be visually identified through a 40x objective on the microscope (Olympus) by their EYFP-fluorescence. Whole-cell clamped recordings were performed on these NG2+ cells using glass capillaries filled with a Cesium Gluconate-based internal solution. The resistance of the recording electrode was 3-6 MΩ, and the NG2+ cells were voltage clamped to a holding potential (Vm) of −60 mV. Voltage-gated sodium currents were activated by applying 20 mV steps ranging from −120 mV to +40 mV with rectangular stimuli lasting 199 ms by use of an AxoPatch 200B amplifier (Molecular Devices, California, USA). Data were acquired with PClamp11 software (Molecular Devices, California, USA), lowpass Bessel filtered at 1 kHz and sampled at 50 kHz. For Detailed description on the electrophysiology experiments and on data analysis are provided in the SI.

### Statistics

All samples represent biological replicates. Sample sizes are indicated in the figure (Fig) legends. Values are expressed as mean ± SEM. Data were checked for normal distribution using the Kolmogorov-Smirnov test. Unpaired two-tailed Student’s t test (normally distributed) or Mann– Whitney U-test (not normally distributed) were used to compare sets of data obtained from two independent groups of animals. Pearson’s correlation coefficient was used to measure linear correlation between two sets of data. P values are reported in Fig legends, with P < 0.05 considered statistically significant. All data were analyzed using Prism version 8.3 (GraphPad Software).

## Results

### Changes of the NG2+ cell transcriptional signatures in response to ELS: impact of glucocorticoids

We used a very well-established model for memory deficit, the ELS (40,41,45). Similar to previous published studies, we detected increased plasma CORT concentrations (Fig 2A) and a decrease in body weight (Supplementary Table 1) in pups exposed to ELS. The CORT concentrations varied highly across ELS litters (Fig 2B) despite identical experimental conditions. We therefore stratified the ELS litters based on the CORT concentration showed by the control into ELS with low CORT concentration (≤5 ng/mL) and ELS with high CORT concentration (≥ 5 ng/mL). To further investigate whether ELS specifically targets NG2+ cells in brain areas implicated in the behavioral changes seen after ELS exposure, we isolated NG2+ cells from the hippocampus of pups (P9, directly after ELS or respective control pups) and performed bulk RNA-seq followed by differential gene expression (DGE) analysis. When comparing control samples vs all ELS samples, we found only 15 significantly differentially expressed genes (DEGs, 8 upregulated, 7 downregulated, Supplementary Table 2). However, since the impact of ELS in terms of stress-induced increase in plasma CORT concentration varied highly between ELS litters, we divided the samples based on the control level into low-CORT ELS (≤ 5 ng/mL), high-CORT ELS (≥5 ng/mL) and controls, which always had plasma CORT concentrations below 5 ng/mL (Fig 2B). Comparison of the RNA-seq data using this stratification revealed 54 DEGs (19 upregulated, 35 downregulated) between controls versus high-CORT ELS samples (Fig 2C and Table 1). No significant DEGs could be detected for control vs low-CORT ELS and low-CORT ELS vs high-CORT ELS. Further analysis revealed that the expression of some candidate DEGs (e.g *Scn7a*, *Nwd1*, *Grb10*) across samples strongly correlated with the plasma CORT concentrations (values of r ≥ 0.8) (Fig 2C), whereas some candidates only showed very weak correlation with the stress hormone (r ≤0.3) (e.g *Agpat5, Fbf1, Ncam1*). A significant functional enrichment of our candidate genes in terms of overrepresented molecular pathways could not be found. However, several candidate genes are involved in lipid metabolic processes (*Pnlip*, *Enpp6*, *Slc27a1*, *Agpat5*, *Fa2h*, *Neu4*).

**Table 1:** DE candidate genes (control vs high CORT comparison) sorted by significance. The table shows official gene symbols, effect size (log2-fold change) and significance (adjusted p-value) of differential expression. For the correlation analysis of CORT values and gene’s expression values across samples the table presents Pearson correlation coefficients and associated p-values. “PC12” and “Hippocampus” indicate whether a gene has been detected to differentially bind in CORT-associated ChIP-seq studies (see text).

| gene symbol | log2FoldChange | padj | correlation | P-value | PC12 | Hippocampus |
| --- | --- | --- | --- | --- | --- | --- |
| *Scn7a* | 1.07 | 2.51E-13 | 0.82 | 0.00 | x |  |
| *Bmp4* | -0.78 | 4.48E-07 | -0.61 | 0.00 |  |  |
| *Nwd1* | 0.73 | 2.80E-06 | 0.91 | 0.00 |  |  |
| *Moxd1* | 0.73 | 1.96E-05 | 0.64 | 0.00 |  |  |
| *Shisa9* | -0.56 | 1.96E-05 | -0.56 | 0.01 |  |  |
| *Grb10* | 0.61 | 1.97E-05 | 0.83 | 0.00 | x |  |
| *Vit* | 0.72 | 3.88E-05 | 0.60 | 0.00 |  |  |
| *Tnfaip8* | 0.63 | 1.29E-03 | 0.75 | 0.00 |  |  |
| *Gm44667* | 0.48 | 1.57E-03 | 0.41 | 0.06 |  |  |
| *Dock9* | -0.43 | 1.57E-03 | -0.50 | 0.02 |  | x |
| *Galnt6* | 0.62 | 3.13E-03 | 0.24 | 0.27 |  |  |
| *Peg3* | 0.31 | 3.80E-03 | 0.53 | 0.01 |  |  |
| *Msrb3* | -0.58 | 3.80E-03 | -0.61 | 0.00 |  |  |
| *Fa2h* | -0.54 | 4.12E-03 | -0.44 | 0.04 | x | x |
| *Slc16a12* | -0.57 | 0.01 | -0.55 | 0.01 |  |  |
| *Slc27a1* | 0.41 | 0.01 | 0.76 | 0.00 |  |  |
| *Faim2* | -0.52 | 0.01 | -0.59 | 0.01 |  |  |
| *Mmp14* | 0.48 | 0.01 | 0.68 | 0.00 |  |  |
| *Cd9* | -0.47 | 0.01 | -0.59 | 0.00 |  |  |
| *Lbh* | -0.45 | 0.01 | -0.62 | 0.00 |  |  |
| *Sirt2* | -0.43 | 0.01 | -0.56 | 0.01 |  |  |
| *Igf2bp2* | 0.52 | 0.01 | 0.48 | 0.02 |  |  |
| *Slc5a7* | -0.5 | 0.01 | -0.41 | 0.05 | x |  |
| *Lrrn1* | -0.39 | 0.02 | -0.56 | 0.00 |  |  |
| *Psenen* | -0.47 | 0.02 | -0.66 | 0.00 |  |  |
| *Tle2* | 0.46 | 0.02 | 0.29 | 0.19 |  |  |
| *Bmp3* | -0.53 | 0.02 | -0.38 | 0.08 |  |  |
| *Cfdp1* | -0.35 | 0.02 | -0.69 | 0.00 |  | x |
| *Lrtm2* | -0.48 | 0.02 | -0.66 | 0.00 |  |  |
| *C1ql1* | -0.52 | 0.02 | -0.57 | 0.00 |  |  |
| *Agpat5* | -0.24 | 0.02 | -0.44 | 0.04 |  |  |
| *Btd* | -0.41 | 0.02 | -0.53 | 0.01 |  | x |
| *Dpm2* | -0.43 | 0.02 | -0.64 | 0.00 |  |  |

**Fig 2:**
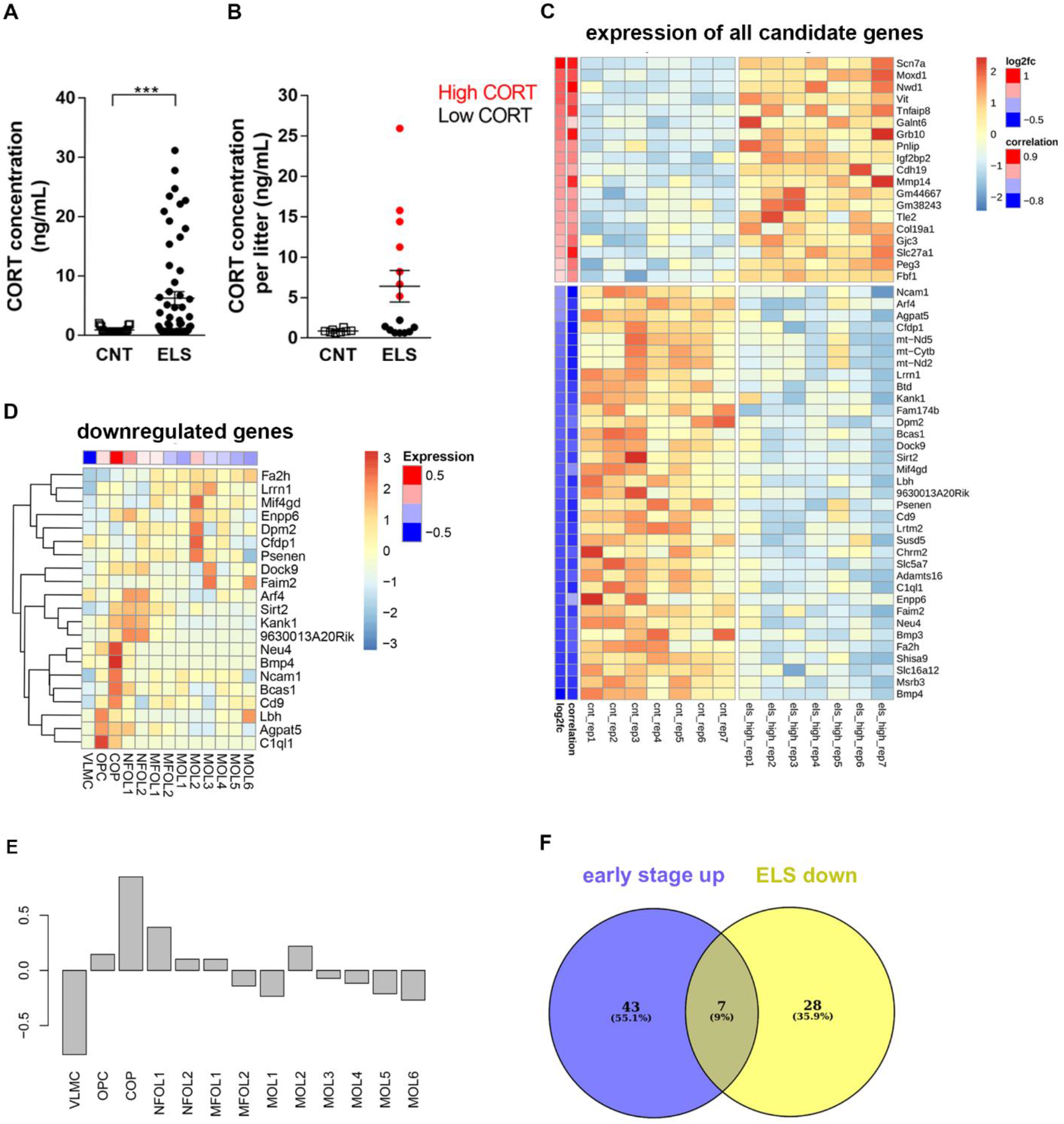
Impact of individual increase in circulating plasma CORT concentrations on ELS-induced transcriptional changes in NG2+ cells and stage-wise expression of ELS-downregulated genes in oligodendrocyte lineage. (A) Graph showing the ELS-induced increase in plasma CORT based on individual values of each animal at P9. Two-tailed Mann-Whitney Test, U= 442, p=0.0009, n= 27 CNT and n= 57 ELS. (B) Averaging the individual plasma CORT concentrations per litter, this graph illustrates the heterogeneity of the CORT response across litters following ELS with the stratification in high CORT and low CORT concentration. Two-tailed Mann-Whitney Test, U= 30, p= 0.0553, n= 8 CNT litters and n= 15 ELS litters. *p<0.05 ***p< 0.001. (C) Heatmap representing DEGs between control and high-CORT litters. In total, 54 genes were significantly upregulated (19) or downregulated (35, |log2fc| > 0.2, padj < 0.05). The heatmap shows scaled expression values for each gene across all relevant samples, whereby higher/lower expression is indicated by red/blue color. The magnitude of change in expression (log2fc) is indicated on the left-hand side of the plot, next to this the Pearson correlation coefficients of gene expression with CORT level in respective samples are shown. (D) The heatmap represents centered and scaled gene expression (mean subtracted, divided by standard deviation) of the ELS-downregulated DEG in each stage of the oligodendrocyte lineage derived from the data available in (46). These values reside between ~-3 and +3 values. The cells on top of the heatmap represented sums of scaled expression values per each cell stage in the lineage. Range −0.5 to +0.5. Abbreviation: newly formed oligodendrocytes (NFOL1 and NFOL2); myelin-forming oligodendrocytes (MFOL1 and MFOL2; mature oligodendrocytes (MOL1 to MOL6). (E) The Panel represents with a bar plot the same values represented in the cells on top of the heatmap. The left part of the heat map shows the clustering dendrogram of genes and represents "patterns" of similar expression, to which the genes are also grouped. (F) The Venn diagram indicates the overlap of genes which are early-stage indicator genes as identified in single-cell study ((46), left-hand side) and downregulated genes in ELS animals (right-hand side).

Since we found a strong correlation between the ELS-induced plasma CORT concentrations and some of our DEGs (Fig 2C and Table 1), we wanted to understand whether the identified DEGs are glucocorticoid responsive genes. We therefore took advantages of already existing data on genomic binding sites of glucocorticoid receptor (GR) in neuronal PC12 cells (47) and hippocampi (48) of rats treated with dexamethasone (DEX), a synthetic glucocorticoid receptor antagonist (49). Overlap analysis of genes binding due to DEX treatment in PC12 cells (918) and our 54 candidate DEGs revealed a significant enrichment represented by 5 genes: *Scn7a*, *Grb10*, *Fa2h*, *Slc5a7* and *Chrm2* (Fisher enrichment fold 2.79, p-value 0.0416; see Table 1 for overlapping genes). Overlap with the rat hippocampus study showed no significant enrichment represented by 5 genes (5 out of 1828/54 candidates, 1.34 fold/p=0.431, see Table 1).

### ELS shifts the transcriptome of NG2+ cells towards a more mature cell lineage

Since NG2+ cells can maturate into oligodendrocytes, we wondered whether the ELS induced change in their transcriptional signature could reflect a shift towards a different stage in the oligodendrocyte lineage. We therefore performed a comparative analysis between our 54 candidate DEGs and previously published transcriptional signatures characterizing six maturation stages (13 substages) of the mouse oligodendrocyte lineage (46). We could show that DEGs downregulated in the ELS samples tended to be higher expressed in early stages of the oligodendrocyte lineage (Fig 2D and E). This tendency was further confirmed by a significant enrichment of 7 out of the 35 downregulated-DEGs and the 50 early-stage indicator genes identified in the single-cell study (46) describing transcriptional signatures for vascular and leptomeningeal cells (VLMC), OPC and differentiation-committed oligodendrocyte precursors (COP) (7 out of 35/50, hypergeometric test p-value=2.013*10−12, as shown in Fig 2F). The expression levels of these seven genes in CNT and ELS samples can be found in Supplementary Fig 1A. We found that the up-regulated DEGs in the ELS samples were not significantly increased or decreased in any of the oligodendroglia lineage stage (Supplementary Fig 1B and C). No other combinations (upregulated DEGs vs early-stage, upregulated DEGs vs late-stage, downregulated DEGs vs late-stage) showed an overlap.

### Long lasting effects of ELS on adult hippocampal NG2+ cell transcriptome and cognitive performance: persistent upregulation of *Scn7a*

To further evaluate the functional impact of the observed transcriptional changes on cognitive integrity, we examined the behavioral outcome 4-5 months after ELS exposure using two behavioral tests: the novel object recognition test (NORT, assessing hippocampus-dependent cognitive performance) and the open field test (OF, investigating locomotor activity). We showed an impaired cognitive performance in the NORT and increased locomotor activity in the OF (Fig 3B and 3C) in ELS exposed mice. We did not detect any difference in the basal plasma CORT concentrations in adult animals previously exposed to ELS compared to control animals (Fig 3A).

**Fig 3:**
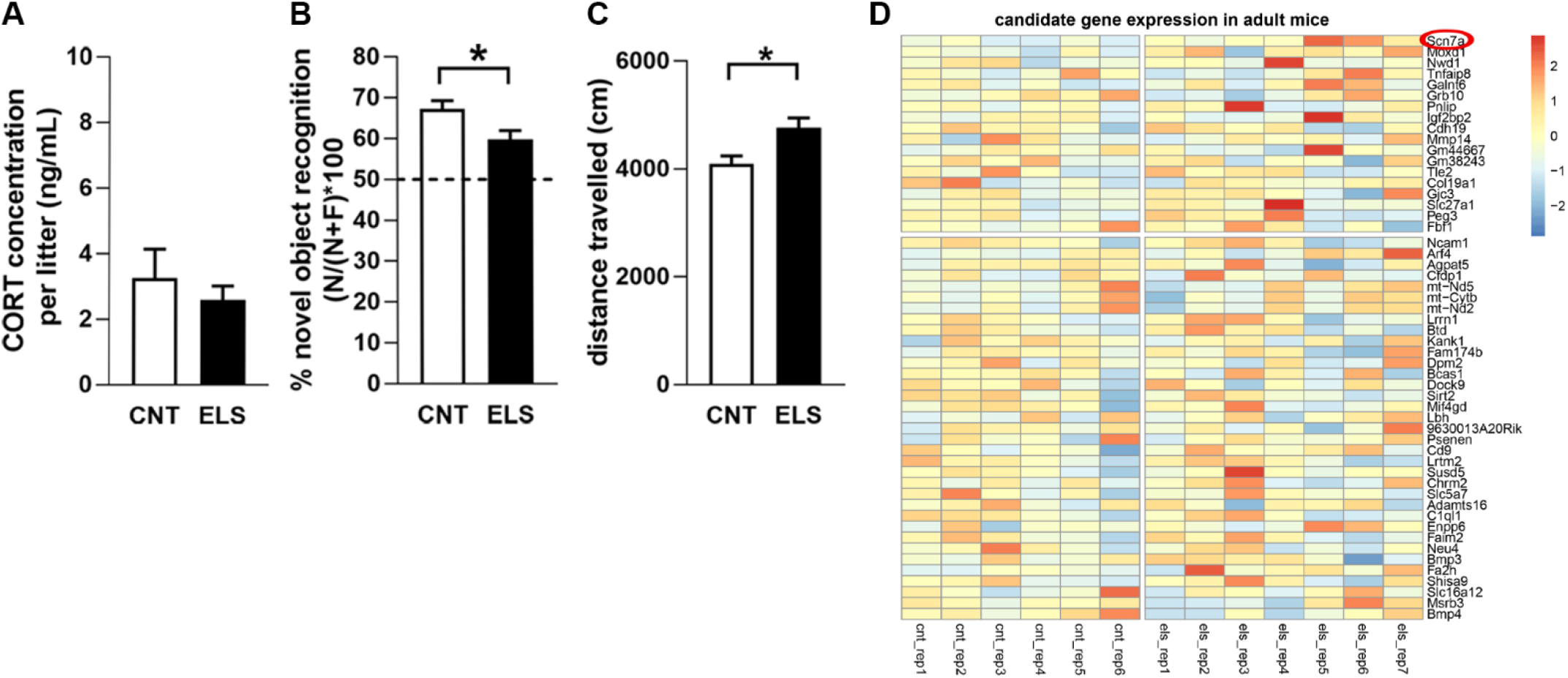
ELS-induced impairment of cognitive performance in adult animals and concomitant upregulation of Scn7a. (A) Graph showing the plasma CORT concentrations in adult animals calculated per litter. Unpaired Student’s t-test t(16)= 0.7235, p= 0.4798, n= 8-11 litter per group. (B) Graph showing the % of novel object recognition of adult animals calculated per litter. Unpaired Student’s t-test t(18)= 2.416, p= 0.0266, n= 8-12 litter per group. (C) Graph showing the distance travelled in the OF of adult animals calculated per litter. Unpaired Student’s t-test t(18)= 2.566, p=0.0194, n= 8-12 litter per group (D) *Heatmap representing the DEGs derived from the P9 analysis (from Fig 3) re-analyzed at adult stage. The heatmap shows scaled expression values for each gene across all relevant samples, whereby higher/lower expression is indicated by red/blue color. Only one gene*, *Vit, was below detection limit (<5 reads in all samples) and was therefore omitted from analysis. Only Scn7a remained up-regulated after performing targeted t-test.*

In order to understand whether the ELS-associated alterations in the transcriptome of NG2+ cells isolated from P9 hippocampi are long lasting and might explain the accelerated cognitive decline, we performed an additional series of NG2+ cell transcriptional profiling on those cognitively impaired, 8-months old animals. At the transcriptome-wide level, we found no DEGs between control and mice previously exposed to ELS, which points to a relative normalization of the NG2 transcriptome profile in the adult stage. However, when choosing a candidate-driven approach and performing a targeted t-test of the same DEGs that had been identified in ELS pups, we found that one DEG (*Scn7a*) was significantly upregulated also in the adult animals (target-t-test p= 0.01969, t (11) = 2.8925, Fig 3D).

### ELS modulates voltage-sensitive sodium currents in hippocampal NG2+ cells

Since *Scn7a* encodes for subunits of sodium channels and it has been previously identified in NG2+ cells (50–53), we further tested the hypothesis that ELS could change the electrical properties, thereby modulating the activity, of hippocampal NG2+ cells. To this end, we investigated the impact of ELS on the density of Voltage-Gated sodium (Nav) channels by the use of whole-cell patch-clamp electrophysiological recordings in heterozygous NG2-EYFP knock-in pups (43, 51) (Fig 4A). As expected, and consistent with our data from the C57Bl/6J strain, the NG2-EYFP line (+/-) mouse was responsive to ELS and showed the characteristic ELS-induced increase in plasma CORT concentrations as well as a decrease in body weight (Supplementary Table 1). In addition, electrophysiological recordings from NG2+ cells, identified through EYFP-expression in hippocampal slices, revealed an increased density of Nav mediated sodium currents in the ELS condition, with a peak between ELS and control conditions at a holding potential of +20 mV (Fig 4B and C).

**Fig 4:**
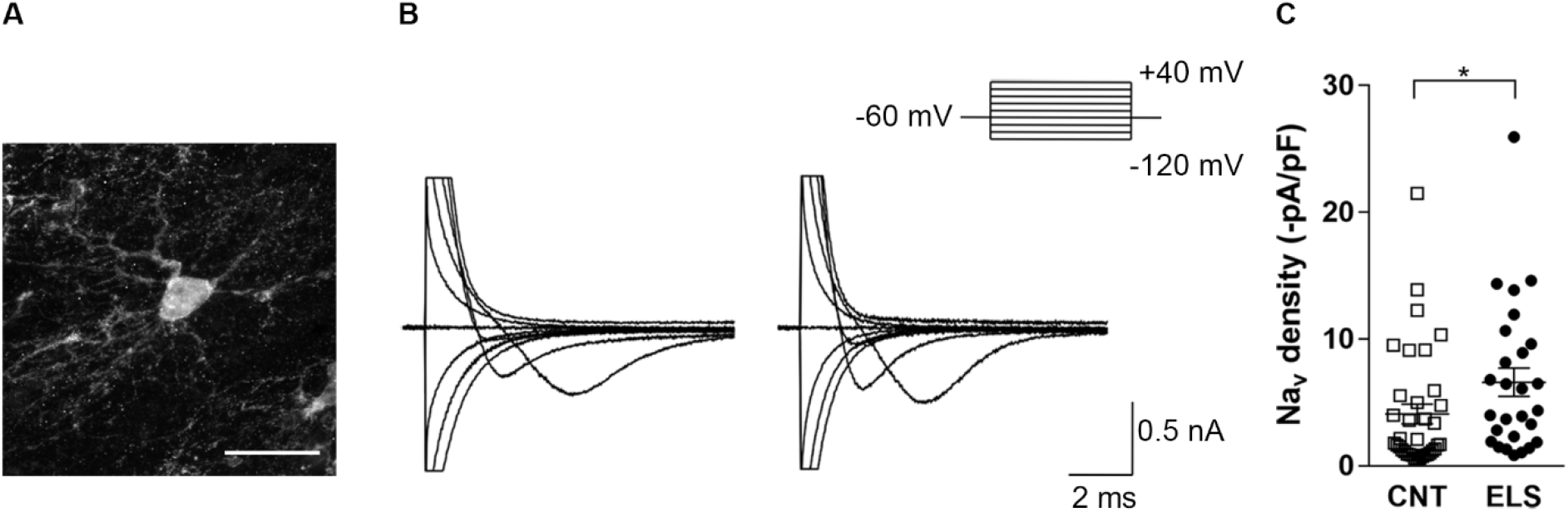
ELS altered the current density of Nav channels in hippocampal NG2+ cells. (A) Representative NG2+ cell. Cells were selected by their EYFP expression (NG2-EYFP knock-in mice) and their characteristic morphology. Scale bar 20 μm. (B) Representative voltage traces of voltage sensitive sodium currents in NG2+ cells from CNT (left) and ELS (right) P9 pups at different holding potentials. (C) Summary plot of mean density of Nav activated sodium currents in control and ELS P9 pups. Two-tailed Mann-Whitney Test, U= 305, p= 0.0115. Data are expressed as mean ± SEM. Unpaired t-test. * p<0.05 vs CNT (30-36 cells in 12 animals per group).

## Discussion

In this study, we have defined a stringent functional link between the impact of early life adversity on cognitive integrity and hippocampal NG2+ glia as a putative mediator of CNS homeostasis. We employed a mouse model of ELS to show that ELS specifically targets the transcriptional signature of hippocampal NG2+ cells, and that changes in the NG2 transcriptome largely depend on the ELS-induced increase in circulating plasma CORT. Through integration of our transcriptome data with stage-specific clusters of marker genes in the oligodendrocyte lineage, we provided first evidence that ELS is able to shift the molecular profiles of NG2+ cells towards a more mature stage in the lineage. The most strongly ELS-induced candidate gene, *Scn7a*, continued to be persistently increased until adulthood. Since *Scn7a* encodes for a subunit of sodium channels, we finally confirmed the functional relevance of the transcriptional change for one player mediating the hippocampal network activity (54) by electrophysiological recordings showing an increase in the density of Nav activated sodium currents after ELS.

In line with previous studies (41, 55), we confirmed a strong impact of ELS on circulating CORT plasma concentrations in P9 pups. The plasma CORT concentrations showed considerable variability between litters under strictly controlled and identical experimental conditions. Despite the large number of studies using ELS as a translationally relevant animal model, individuality in the ELS outcome is a so far neglected aspect. Usually, an increase in plasma CORT concentrations is described as a hallmark of ELS; still the specific role of CORT in shaping the long lasting negative effects of ELS on memory and cognitive function is not finally solved (56, 57). It may thus be that the heterogeneity in CORT response across litters indicates a variance in model efficiency among individuals, and that considering ELS-subgroups (i.e. animals resilient or susceptible to the early adverse manipulation) could be interesting to consider in future studies to enhance the translational value.

Our study shows, for the first time, that the highly dynamic population of NG2+ glia is a specific target of early life adversity. NG2+ glia have repeatedly been described to quickly respond to different types of challenges or insults to the adult CNS (58, 59). The fact that the time window of ELS exposure critically hits the period during which NG2+ cells reach their peak density (first postnatal week) provides a strong scientific rationale to investigate the specific role of NG2+ glia in modulating ELS-associated aversive outcomes. ELS-induced changes in the transcriptional signature of NG2+ cells correlated with the plasma concentrations of the stress hormone CORT. This link between glucocorticoids and NG2 transcriptome changes after ELS was further strengthened by additional bioinformatic analyses: when integrating our transcriptome data with publicly available datasets on genomic binding sites of GR in neuronal PC12 cells and hippocampus of rats treated with DEX (47, 48), we identified several overlapping candidates. Thus, ELS-induced activation of the hypothalamus-pituitary-adrenocortical system, which exposes the developing brain to an excess of glucocorticoids seems to play a crucial role major in shifting gene expression, with possible lasting consequences on NG2+ cell function and, presumably, brain homeostasis. Interestingly, many of the downregulated DEGs overlapped with early-stage marker genes in the oligodendrocyte lineage, which suggests an accelerated maturation of NG2+ cells upon excess glucocorticoid exposure. The significance of this finding should be viewed from the fact that the NG2 population is a prototypical cellular target for stress signals, (34, 60) as NG2+ cells express both GR and mineralocorticoid receptors (MR) throughout the maturational stages of the lineage, from OPCS to mature oligodendrocytes (30). Moreover, CORT treatment has been shown to influence oligodendrogliogenesis in the hippocampus (31–33), which may in turn affect neurotransmission, alter neuronal function and ultimately behavior (also discussed in (32)). Thus, it is tempting to speculate that early adversity, by altering the transcriptome pattern of OPCs may lead to changes in the communication between OPCs and neurons. ELS is able to override the protective SHRP, thus putting the various CNS functions that NG2+ cells homeostatically support immediately at risk. Since NG2+ cells are talking back to the neuronal network (27), such changes could alter neuronal function and ultimately lead to long lasting effects on adult emotional behavior and cognitive performance as observed in ELS animals. In support of this hypothesis, Teissier and colleagues found that maternal separation – i.e. different from ELS - caused an increase in mature oligodendrocytes in stressed pups and a consequent decreased in the number of OPCs in adult animals. Comparable results on the OPC/oligodendrocyte ratio were obtained when medial prefrontal cortex neuron excitability was reduced using designed receptors exclusively activated by designed drug (DREADD) technique during the first 2 postnatal weeks in control pups which mimicked ELS-like behavioral alterations later in adulthood (17). Hence, it may well be that ELS, by accelerating the maturation of the OPCs, alters the communication between NG2+ cells and neurons. Recent literature showing that EA results in precocious differentiation of OPCs and neural maturation in both mice and humans (18, 61) indeed supports this idea. However, whether an acceleration in OPC differentiation can cause changes in NG2+ cell-neuron-communication and finally lead to negative behavioral outcomes is unclear, but is indeed likely, as with maturation the expression of NG2 is lost concomitant with the loss of synaptic contact between OPCs and neurons (62, 63).

Considering the remarkable potential of NG2+ cells to homeostatically modulate brain function, ELS-induced upregulation of *Scn7a* in NG2+ cells is an intriguing finding. This gene encodes for a subunit of sodium channels widely expressed by OPCs throughout their lifespan. The channels are fundamental in the transduction of neuronal input onto OPCs (51). Given that *Scn7a* remained persistently upregulated until adulthood, was highly correlated with the concentration of circulating CORT at P9 and differentially bound in ChIP-seq studies targeting GR-responsive transcripts, it is not unlikely that the *Scn7a* is involved in modulating ELS-induced behavioral changes observed in our adult mice. So far, evidence for an involvement of Scn7a in psychiatric disease phenotypes is limited to a recent genome-wide association study showing that a single nucleotide polymorphism in the *Scn7a* gene is associated with response to treatment in depressed patients (64). Data on the exact role of Nav channels in NG2+ cells both under physiological as well as disease-predisposing conditions such as early life stress are still lacking Patch-clamp recordings have confirmed the expression of functional Nav channels in NG2+ cells (65–69). Interestingly, the expression of Nav channels in NG2+ cells follows specific temporal dynamics and changes depend on the developmental stage (65), with higher Nav channel density in the NG2+ proliferative stage (51) but decreased density upon maturation (50, 66, 69). Under physiological conditions, the expression of Nav channels is relevant for OPC-neuron communication. It is known that action potential activity can lead to vesicular glutamate release and consequent activation of ionotropic glutamate receptors in NG2+ cells (70), and that NG2+ cells respond to this type of neurotransmitter release (67). Consequently, Nav channel-mediated electrical activity may serve as a signal between unmyelinated axonal sections and OPCs that are ready to differentiate into oligodendrocytes and are in turn capable of myelinating these axonal targets (65, 67, 71). This kind of electrical activity may thus be crucial in guiding the differentiation of OPCs. Interestingly Nav channels are sensitive to the cellular milieu and alterations in the constituents of the medium are sufficient to alter the expression of functional ion channels (51). Considering that ELS and an excess of glucocorticoid hormones might exert a plethora of cellular and molecular effects it is tempting to speculate that ELS, most likely via an increase in circulating glucocorticoids, enhances Nav density in NG2+ cells. This increase in Nav density may alter the responsiveness of NG2+ cells to neurons with long-lasting negative outcomes on brain function and complex behavioral phenotypes. Thus, is not surprising to see that a recent single cell RNA-seq study from autopsy brain from depressed patients revealed gene alterations specifically in OPCs and excitatory neurons with possible implications for OPC-to-neuron communication (72). Interestingly, in disease conditions, Nav channel expression switches were observed in other non-neuronal cells such as reactive astrocytes, which displayed upregulated expression of Nav channels in acute and chronic multiple sclerosis plaques as well as in tissues surrounding cerebrovascular accidents and in brain tumors from autopsy human brains (65, 73).

In conclusion, our study substantially advances our knowledge about the molecular and cellular mechanisms through which adverse early experiences can be translated into lasting changes of brain function which, later in life, might predispose to the development of mental disorders. We are confident that our data will lay the basis for future studies investigating specific effects of CORT on the maturation of NG2+ cells and examining the effects of stress on NG2+ cell-neuron-communication.

Specifically, restoration of an ELS-induced dysfunction in NG2+ cell-neuron communication could represent a conceptually novel and promising therapeutic avenue to explore in the future.

## Acknowledgment

This work was supported by a DFF Postdoctoral grant from the Danish Research Council (DFF-5053-00103) and from the Dr.phil. Ragna Rask-Nielsen Grundforskningsfond both to GT, and from the German Research Foundation to TM (CRC 1080, TP CO_2_), from the German Research Foundation to JT and TM (MI 452/6-1) and from the German Foundation to JT (SFB CRC 128, TP B07). The support by the IMB Genomics Core Facility in Mainz, the use of its NextSeq500 (INST 247/870-1 FUGG) and the support by the IMB Microscopy Core Facility are gratefully acknowledged. We are very grateful to Dr Inge Sillaber (Genevention), Dr Michael Van der Kooij and Dr Filippo Calzolari for critical discussion and to Franziska Mey, Anna-Lena Schlegelmilch, Annika Hasch, Verena Opitz, Julia Deuster and Jennifer Klüpfer for technical support.

## Conflict of interests

GW reported having received lecture/consultancy fees from H. Lundbeck A/S, Servier SA, Astra Zeneca AB, Eli Lilly A/S, Sun Pharma Pty Ltd, Pfizer Inc, Shire A/S, HB Pharma A/S, Arla Foods A.m. b.A., Alkermes Inc, and Mundipharma International Ltd., Janssen AB. All other authors report no biomedical financial interests or potential conflicts of interest. This manuscript has been posted on a preprint server.

