## Supplementary materials and tables for "Early life stress programming of NG2+ glia transcriptome alters functional properties of voltage gated sodium (Nav) channels and cognitive performance"

**Supplementary information**

### Materials and Methods

#### Animals

Adult female and male C57bL/6J mice (10 weeks) were obtained from Janvier Labs (France). Mice were single housed with food and water ad libitum in an air-conditioned (temperature=22 ± 2 °C, relative humidity=50 ± 5%) housing room with 12 h/12 h light-dark cycle (lights on at 07:00 a.m.). Mice were habituated to the new environment for at least 1 week before the starting of the experiment. The breeding pairs consisted of female and male C57BL/6J. After mating pregnant dams were single housed and put in a cabinet with controlled ventilation, temperature and light cycle until the end of the experiment. From postnatal day (P) 2 to P9 the dams and litter were exposed to early life stress procedure (ELS) or control condition. For electrophysiological experiments C57bL/6N mice were bred to homozygous knock-in NG2 enhanced yellow fluorescent protein (NG2-EYFP) mice, where the expression of the reporter gene is regulated according to the endogenous NG2 promoter, allowing unbiased sampling of this population (1), to obtain heterozygous NG2-YFP pups. All experiments were performed in accordance with the European directive 2010/63/EU for animal experiments and were approved by the local authorities (Animal Protection Committee of the State Government, Landesuntersuchungsamt Rheinland-Pfalz, Koblenz, Germany).

#### Early life stress protocol

ELS was performed as previously described (2, 3). On P2, dams with 5-6 pups (of which at least one female) were randomly subjected to ELS or assigned to the control group from P2 to P9.

The two groups were housed in different shelves of the same cabinet. The dams subjected to ELS were housed in cages with limited bedding and nesting materials (half square of a Nestlet, # 14010 Plexx) on an aluminum mesh platform as for Rice et al., 2008 (McNichols Co) at least 1.5 cm above the cage floor. Control litters were kept under standard conditions and provided with sufficient bedding and nesting materials (2 Nestlets) and few little pieces of the old nesting materials were kept.  All animals were monitored but left undisturbed until the end of the experiment. For the acute effect of ELS, in the morning of P9 the male pups were weighed and euthanized. For the long-lasting effect of ELS at P9 animals were weighed and only 3 males per litter were kept and housed again under standard conditions until weaning at P21. After weaning the male offspring were kept group housed until 8 weeks. At week 9-10 the animals were single housed, behavioral phenotype tested at 4-5 months and mice were euthanized at 8 months. For a schematic overview of the different experimental schedules see Fig 1.

**Measurement of plasma CORT**

Concentrations of circulating corticosterone were measured from trunk blood plasma (P9) or tail cut (adult animals) by Corticosterone ELISA Kit (Enzo Life Sciences, #Cat.No. ADI-900-0979) according to the manufacturer's instructions. The blood was collected in an EDTA tube and centrifuged for 10 min at 4 °C and 10 000 rpm.

**Behavioral tests**

All behavioral experiments were conducted between 09:00 and 14:00 by experimenters blinded to the treatment groups. All the animals were randomized per condition. The videos of all behavioral tests were scored manually by an experimenter blinded to the animals' treatment using The Observer XT12 software (Noldus Information Technology). Between each test session, setups and arenas were cleaned with a 5% EtOH solution and dried with tissues. Behavioral testing occurred in custom-made sound-attenuating boxes under constant light conditions (37 lx).

**Open field (OF)**

The OF consisted of a rectangular arena (45x45x41 cm). Mice were introduced near the wall of the arena and allowed to explore for 10 min. We measured locomotor activity during the OF test.

**Novel object recognition test (NORT)**

Mice were habituated to the arena for two days (during the open field session and on the following day for 20 minutes). On the third day the NORT test was performed. During the habituation phase of the test animals were placed in the arena containing two similar objects at equidistant location and allowed to freely explore for 10 min (familiarization). After a 24h inter-trial interval in the animals' home cage, mice were replaced for 5 min into the arena, now containing one familiar object and a novel object (similar in size but different in shape, material, texture and contrast). The arena and objects were cleaned with 5% EtOH in between trials. Object exploration was scored as direct interaction with the object such as sniffing or touching the object with the nose or forepaws. Trials in which total exploration time was <5s were considered insufficient and removed from the analysis. The time exploring the novel object divided by the total duration of exploration was taken as novel object recognition index (expressed in %).

**Isolation of NG2+ cells by magnetic cell sorting**

NG2+ cells were isolated using the NTDK-P Kit (Miltenyi Biotec) according to previously described procedures (4, 5), where Magnetic isolation (MACS) was performed with anti-NG2 antibody conjugated with magnetic beads (Miltenyi Biotec). For the P9 time-point, each sample was a pool of the hippocampi from the male pups of the entire litter; n represents litter number (control n = 7, ELS low CORT n= 8, ELS high CORT n= 7, randomized in 3 batches). For the 8-month time-point each sample was a pool of the hippocampi of 3-4 males per sample (control n = 6, ELS n = 7 randomized in 3 batches).

**RNA extraction and Next-Generation Sequencing (NGS)**

After NG2 sorting, cells were pelleted and RNA was extracted using RNeasy Micro Kit (QIAGEN) according to manufacturer's instructions. NGS library prep was performed with NuGen Ovation SoLo RNA-Sequencing (RNA-seq) System following NuGen`s standard protocol (M01406v2). For the first experiment (at P9) libraries were prepared with a starting amount of 1 ng and amplified in 14 PCR cycles. Libraries were profiled in a High Sensitivity DNA on a 2100 Bioanalyzer (Agilent technologies) and quantified using the ddPCR Library Quantification Kit for Illumina TruSeq in a QX200 Droplet Digital PCR system (BioRad). All 22 samples were pooled in equimolar ratio and sequenced on 2 NextSeq 500/550 Flowcell, SR for 1x 70 cycles plus 16 cycles for the index read and 5 dark cycles upfront. For the second experiment at 8 months NGS library prep was performed as above and libraries were prepared with a starting amount of 1.355ng (except sample 8 with 0.768 ng) and amplified in 14 PCR cycles. Libraries were profiled in a High Sensitivity DNA on a 2100 Bioanalyzer (Agilent technologies) and quantified using the Qubit dsDNA HS Assay Kit, in a Qubit 2.0 Fluorometer (Life technologies).

All 13 samples were pooled in equimolar ratio and sequenced on 1 NextSeq 500 Highoutput Flowcell, SR for 1x 75 cycles plus 16 cycles for the index read and 5 dark cycles upfront.

**Bioinformatic analysis**

Raw read files were analyzed with FastQC (version 0.11.8, www.bioinformatics.babraham.ac.uk/projects/fastqc/) and found to be of high quality (data not shown). Reads were mapped to the mouse genome (version GRCm38.p6) using the Star alignment software (version 2.5.3a, (6)) resulting in unique mapping rates >90% with ~30-45 mio mapped reads per sample. Since a low input RNA-seq protocol with a high number of PCR cycles was used for RNA extraction, a deduplication procedure was applied to alignment (BAM) files using a kit manufacturer's python script (NuDUP 2.3.3, https://github.com/nugentechnologies/nudup) and the UMI indices in raw read files (~40% deduplicated read fraction per sample). Transcript quantification was performed using the FeatureCounts software of the Subread package (version 1.5.3,(7)) and GENCODE mouse annotation (version M17,(8)).

Linear and nonlinear dimension reduction for 2-d transcriptome profile representations were performed through PCA and t-SNE implementations in the R analysis environment. Differential gene expression analysis was conducted using the DESeq2 R package (version 1.20.0, (9)). We considered only transcripts with at least 5 mapping reads in one of the corresponding samples. We used DESeq2 with default parameters and the integrated log2-fold change shrinkage method 'normal'. Significantly differentially expressed genes were identified at adjusted p-value cutoff padj<=0.05. Correlation of gene expression and cort values across samples was performed in R using the cor.test function with Pearson correlation using complete observations only. Pathway and transcription factor enrichment were calculated using the

DAVID and Enrichr online tools (david.ncifcrf.gov, version 6.8, (10) and amp.pharm.mssm.edu/Enrichr/, accessed June 2019, (11)). To identify GR responsive genes, DE genes were overlapped with binding candidates in two rat PC12/hippocampus ChIP-seq studies (12, 13) and Fisher exact tests were performed to assess enrichment scores.

For analysis of cell stage-specific expression of candidate genes and potential cell type differentiation in ELS animals the data from a single-cell RNA sequencing study (14) was analyzed and compared to bulk RNA seq data obtained in this study. To match the brain region under investigation, only cells from the dentate gyrus and hippocampus CA1 were considered. Cell stage-specific expression of candidate genes was calculated by averaging normalized expression values in cells corresponding to stage and cell type annotation as available in the published dataset (GEO accession number GSE75330). Only genes with expression values above a noise threshold (normalized count >= 1) were considered. To determine overlap enrichment of stage-specific genes and ELS regulated genes we calculated hypergeometric test values for four sets according to up- or downregulated ELS genes overlapped with early-stage or late-stage marker genes as obtained from the supplementary material of (14). Expression of stage-specific marker genes in bulk RNA data was determined using the marker gene list published in (14).

**Electrophysiology**

Hippocampal NG2-cells were electrophysiologically characterized at the age of P9-P10 (n=11 (control), n=11 (ELS)) by use of heterozygous knock-in mice expressing the enhanced yellow fluorescent protein EYFP in NG2 cells (NG2-EYFP+/- mice, (1)). After decapitation the brains were quickly removed and put into ice-cold 95% $O_2$/5% $CO_2$ oxygenated ACSF (in mM: 125

NaCl, 3 KCl, 2.5 $CaCl_2$, 1.3 $MgSO_4$, 1.25 $NaH_2PO_4$, 13 Glucose, 26 $NaHCO_3$). Next, the tissue was horizontally cut by use of a vibratome (VT 1200S, LEICA, Germany) to generate hippocampal brain slices with a thickness of 300 µm. The slices were kept in oxygenated ACSF at room temperature for at least 1 hour for recovery. Then they were transferred into a submerged-type recording chamber mounted on an upright microscope (Olympus BX-51WI) and perfused with oxygenated 95% $O_2$/5% $CO_2$ HEPES-buffered recording solution (in mM: 144 NaCl, 2.5 KCl, 10 HEPES, 1 $NaH_2PO_4$, 2.5 $CaCl_2$, 10 Glucose, 0.1 Glycine, 200 $BaCl_2$; pH 7.4 with NaOH) at room temperature for 10 minutes. NG2+ cells located in the stratum radiatum of area CA1 could be visually identified through a 40x objective on the microscope (Olympus) by their EYFP-fluorescence. Whole-cell clamped recordings were performed on these NG2+ cells using of glass capillaries filled with a Cesium Gluconate-based internal solution containing in mM: 125 CsOH, 125 Gluconic Acid, 5 $CsCl_2$, 10 EGTA, 2 ATP-Na2, 2 $MgCl_2$, 0.4 GTP-Na2, 10 HEPES, 5.4 Biocytin hydrochloride; pH 7.3 with CsOH. The resistance of the recording electrode was 3-6 MΩ, and the NG2+ cells were voltage clamped to a holding potential (Vm) of -60 mV. Voltage-gated sodium currents were activated by applying 20 mV steps ranging from -120 mV to +40 mV with rectangular stimuli lasting 199 ms by use of an AxoPatch 200B amplifier (Molecular Devices, California, USA). Data were acquired with PClamp11 software (Molecular Devices, California, USA), lowpass Bessel filtered at 1 kHz and sampled at 50 kHz. Data were analyzed with Clampfit11 and included into analysis when the leak current was ≤ 500 pA and the change of access resistance (ΔRa) pre- and post-stimulation was ≤ 20%. For data analysis, all voltage signals were subtracted from the signal recorded at -100 mV and the sodium channel density (-pA/pF) is presented at a holding potential of +20 mV. An additional stimulation protocol was performed in some cells in presence of the voltage-gated sodium channel blocker TTX (0.5 µM, bath applied). As expected, those recordings failed to induce any inward currents confirming the origin of our activated voltage signal. All electrophysiological recordings were performed between 09:00 and 14:00 by experimenters blinded to the animal groups.

**Immunohistochemistry**

After recordings, 300 µm thick slices were fixed in 4% paraformaldehyde for 1 h at RT, incubated with sucrose 15% overnight and then left in sucrose 30% until farther processing. Slices were cryo-sectioned at 50 µm thickness and stored in PBS until use. Immunohistochemistry was performed as previously described (15, 16) and slices were incubated with a primary antibody against GFP overnight at 4°C. Representative fluorescence images were acquired with a Leica SP5 (LAS AF software) using a 40x oil (NA 1.3) or a 63x oil (NA 1.4) objective (HC PL APO CS2 UV) as z-stacks with a height of ca. 5 µm. The fluorophore was excited with an Argon laser (488 nm) and the fluorescence was detected with a HyD detector in the photon counting mode (534 -588 nm). Images are presented as a maximum projection along the z-axes. Brightness and contrast adjustments were adapted to the minimum and maximum intensities of the intensity histogram using the open source software Fiji (v1.53).

**Statistics**

All samples represent biological replicates. Sample sizes are indicated in the figure (Fig.) legends. Values are expressed as mean ± SEM. Data were checked for normal distribution using the Kolmogorov-Smirnov test. Unpaired two-tailed Student’s t test (normally distributed) or Mann-Whitney U-test (not normally distributed) were used to compare sets of data obtained from two independent groups of animals. Pearson’s correlation coefficient was used to measure linear correlation between two sets of data. P values are reported in Fig legends, with $P < 0.05$ considered statistically significant. All data were analyzed using Prism version 8.3 (GraphPad Software)

### Supplementary figures, tables and legend

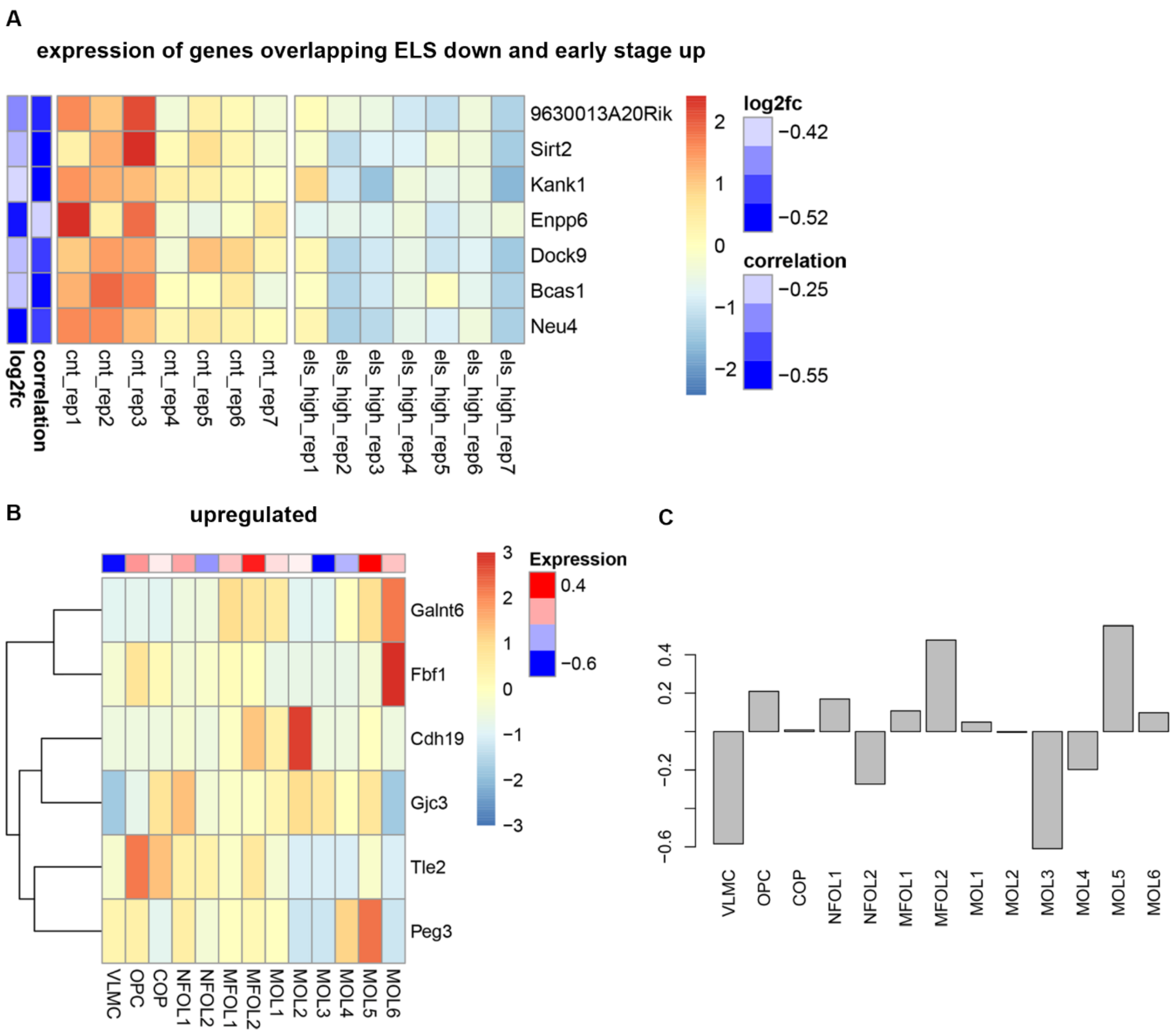


**Supplementary Fig 1: Stage-wise expression of ELS-regulated genes in oligodendrocyte lineage**

(A) Expression patterns in CNT and ELS samples from P9 pups of the7 genes overlapping between our 35 downregulated-DEGs (see Fig 2) and the 50 early-stage (VLMC, OPC, and COP) indicator genes identified in the single-cell study (14). The heatmap (B) represents centered and scaled gene expression (mean subtracted, divided by standard deviation) of the ELS-upregulated DEG in each stage of the oligodendrocyte lineage derived from the data available in (14). These values reside between ~-3 and +3 values. The cells on top of the heatmap represents sums of scaled expression values per each cell stage in the lineage. Range -0.6 to +0.4. (C) The Panel represents with a bar plot the same values represented in the cells on top of the heatmap.

**Supplementary Table 1**

| | Parameter | CNT | ELS | Statistics |
|---|---|---|---|---|
| P9 C57bL/6 pups | Body weight | 4.489 ± 0.2582 | 3.337±0.1387*** | Unpaired Student´ t-test; *t(20)= 4.304, p= 0.0003* |
| Adult C57bL/6 mice | Body weight | 33.09 ± 1.016 | 28.76± 0.4214*** | Unpaired Student´ t-test; *t(18)= 4.465, p= 0.0003* |
| P9 NG2-YFP pups | Body weight | 5.404 ± 0.2408 | 3.510± 0.1258**** | Unpaired Student´ t-test; *t(21)= 7.440, p< 0.0001* |
| P9 NG2-YFP mice | CORT level | 2.199 ± 0.4525 | 6.803 ± 1.441* | Unpaired Student´ t-test; *t(17)= 2.374, p= 0.0296* |

**Supplementary Table 1: Physiological effects of ELS in pup and adult mice**

The table shows the weight of CNT and ELS animals at P9 and adult age and the weight and CORT level at P9 of the NG2-YFP mice exposed to ELS. Data are presented as mean ± SEM; *p< 0.05, ***p< 0.001, ****P< 0.0001.

**Supplementary Table 2**

| # | gene symbol | log2FoldChange | padj |
|---|---|---|---|
| 1 | Scn7a | 0.70 | 0.0001 |
| 2 | Shisa9 | -0.49 | 0.0001 |
| 3 | Bmp4 | -0.57 | 0.0030 |
| 4 | Gm44667 | 0.43 | 0.0035 |
| 5 | Dock9 | -0.37 | 0.0080 |
| 6 | Galnt6 | 0.58 | 0.0142 |
| 7 | Slc5a7 | -0.47 | 0.0193 |
| 8 | Tle2 | 0.45 | 0.0193 |
| 9 | Mif4gd | -0.44 | 0.0193 |
| 10 | Nrarp | -0.46 | 0.0348 |
| 11 | Igf2bp2 | 0.47 | 0.0348 |
| 12 | Fa2h | -0.45 | 0.0348 |
| 13 | Grb10 | 0.43 | 0.0401 |
| 14 | Moxd1 | 0.50 | 0.0413 |
| 15 | Tnfaip8 | 0.49 | 0.0483 |

**Supplementary Table 2: Significantly differentially expressed genes between control and all early life stress samples.**

The table shows official gene symbols, effect size (log2-fold change) and significance (adjusted p-value) of differential expression.
